# Eye gaze is not inversion-proof: A robust, sex-invariant gaze inversion effect

**DOI:** 10.1101/2024.04.10.588791

**Authors:** Daisuke Matsuyoshi, Kana Kuraguchi, Hiroshi Ashida, Katsumi Watanabe

## Abstract

Humans are adept at distinguishing individual faces, yet inversion dramatically impairs this ability. This face inversion effect is remarkably robust across observers, but evidence is mixed as to whether inversion also impairs the perception of facial parts, particularly the eye region. Some studies have shown that featural processing is preserved or even enhanced when faces are inverted, whereas others have reported clear inversion-related impairments in feature-based judgements. These mixed findings may reflect limited statistical power, unbalanced participant sex ratios, and heterogeneous task designs. To address these issues, we examined how strongly face inversion affects sensitivity to gaze direction in a well-powered, sex-balanced sample. A total of 190 participants judged whether the eyes in briefly presented upright or inverted faces were looking directly at them or not. Inversion reliably reduced sensitivity to gaze direction, yielding a medium-to-large effect size. Females showed modestly higher overall sensitivity than males (a small-to-medium effect), whereas the inversion effect was highly similar for females and males. These findings show that brief gaze judgements are not immune to inversion, even in a task that could in principle be based largely on eye-region information. They provide quantitative constraints that models of gaze perception should accommodate, including the largely sex-invariant inversion effect.

## 1. Introduction

Humans are experts at distinguishing and recognizing faces under various conditions. People can detect subtle differences and distinguish between thousands of individuals (Jenkins et al., 2018) even though typical human faces share a strikingly similar overall structure (i.e., two eyes, a nose, and a mouth). However, this ability deteriorates markedly when faces are inverted (Yin, 1969). Turning a face upside-down makes it particularly difficult to recognise, whereas inverting many other types of objects does not affect recognition performance as much.

This face inversion effect (FIE) is exceptionally robust across observers and tasks. It has led researchers to speculate that there are *qualitative* differences between upright and inverted face processing, with a shift from holistic or configural processing for upright faces to more piecemeal or featural processing for inverted faces (Maurer et al., 2002). From this qualitative view, perceiving a face as a coherent whole (rather than as a collection of separate elements) is an idiosyncratic feature that makes face recognition special (Farah et al., 1998; Moscovitch et al., 1997). A key prediction of this view is that, because of the shift in processing mode, the processing of individual features should be comparatively spared when a face is presented upside down. By contrast, more recent studies have suggested that the differences in processing upright and inverted faces are *quantitative* (i.e., a matter of degree in processing efficiency) rather than qualitative (i.e., invoking distinct processing mechanisms) (Gold et al., 2012; Jiang et al., 2006; Matsuyoshi et al., 2015; Sekuler et al., 2004). Sekuler et al. (2004), for example, used a response-classification technique to identify the most discriminative stimulus regions that drove performance and found that observers relied on similar local regions across orientations but extracted information less efficiently from inverted faces. Gold et al. (2012) further showed that face perception is no better than would be predicted by Bayesian integration of information from individual facial features. If observers used a truly holistic face representation, they should exhibit superoptimal performance for whole faces (better than the sum of their parts), yet performance was comparable to the Bayesian prediction derived from features presented in isolation. These findings are difficult to reconcile with a strict qualitative shift and instead support a quantitative view in which similar recognition strategies operate for both upright and inverted faces, albeit less efficiently for the latter. At the same time, the two views are not mutually exclusive, and dissociable neural pathways may still contribute to relatively holistic and more feature-based mechanisms (Matsuyoshi et al., 2015).

The behavioural FIE itself is very strong and has been confirmed by rigorous replications. However, it is less clear whether inversion impairs the perception of individual facial features. In particular, whether sensitivity to information conveyed by the eyes, such as gaze direction, varies with face orientation remains unclear. Several studies have reported that inversion has little to no effect on the ability to distinguish faces that differ only in the shape of individual features (Freire et al., 2000; Le Grand et al., 2001). For example, upright faces with different hairstyles and costumes tend to be perceived as different individuals even though their internal features are identical (Sinha & Poggio, 1996), and composite-face paradigms (Murphy et al., 2017) have shown that inversion can markedly improve the ability to identify individuals based on their internal features (Hole, 1994; Young et al., 1987). Xu and Tanaka (2013) also reported that inversion had little or no effect on discriminating featural and configural differences in the eye region (while severely disrupting the perception of changes in the lower region of the face). These results suggest that featural processing, at least for some aspects of eye or gaze information, is preserved or even enhanced when faces are upside-down. By contrast, numerous studies have reported the opposite pattern, namely that featural processing becomes more difficult when faces are inverted. One prominent example is the Thatcher illusion (Thompson, 1980), in which observers fail to notice local feature changes in an inverted face, even though the changes are obvious in an upright face. Other studies have suggested that isolated features presented without a full face context can still produce inversion effects (Leder et al., 2001; Rakover & Teucher, 1997). Taken together, the literature provides evidence for both relative preservation and clear costs of inversion in processing individual features, making it difficult to reach consensus on how inversion affects feature-level information.

Methodological differences among studies may contribute to these discrepancies. Sample size, participant demographics, study design, and task settings all vary substantially across the literature. In particular, sample size and participant sex composition are critical factors in face processing research. Insufficient statistical power due to small samples can increase the likelihood of false positives and undermine the reliability of findings (Button et al., 2013; Simmons et al., 2011). Moreover, females often outperform males in face processing tasks (Bayliss et al., 2005; Goodman et al., 2012; Heisz et al., 2013; Matsuyoshi et al., 2014; Matsuyoshi & Watanabe, 2021; McClure, 2000), yet studies have not always monitored or balanced the sex ratio unless the sex differences were the primary focus. Previous inconsistencies may therefore partly reflect unaccounted variance in participant demographics. Given the recurring reports of a female advantage and sex as a potential moderator across diverse face processing tasks (Cellerino et al., 2004; Lewin & Herlitz, 2002; Matsuyoshi et al., 2014), balancing the sample for sex and explicitly testing for sex effects may help to obtain more precise estimates of the inversion cost.

In the present study, we examined the gaze inversion effect, namely the impact of face inversion on sensitivity to small deviations from direct gaze. Our primary goal was to obtain a precise estimate of the extent to which face inversion affects sensitivity to gaze direction under tightly controlled psychophysical conditions in a large, sex-balanced sample. Observers judged whether the eyes in briefly presented faces were looking directly at them, with faces shown upright or inverted. The judgement is driven primarily by eye-region cues, although observers may also use broader facial context. It therefore provides an informative test of whether inversion costs are confined to clearly global, configural aspects of faces or also extend to eye-region–based judgements. A second, more exploratory goal was to examine whether the inversion effect on gaze discrimination differs between females and males, building on prior reports suggesting that sex may moderate performance on tasks involving facial cues. This design allowed us to quantify the magnitude of inversion costs on gaze discrimination and to assess whether these costs generalise across sexes, thereby providing quantitative constraints that theories of gaze perception should accommodate.

## 2. Materials and Methods

### 2.1 Participants

We assumed a medium-sized effect of sex on gaze direction discrimination based on studies showing sex differences in face or gaze perception (Bayliss et al., 2005; Goodman et al., 2012; Matsuyoshi et al., 2014; Matsuyoshi & Watanabe, 2021). To achieve a power (1 − *β*) of 0.95 and a medium effect size (Cohen’s *f* = 0.25) while assuming a high correlation between the measures (*r* = 0.7 between upright and inverted gaze processing), a sample of 180 (90 for each group) is required to detect a significant difference (*α* = 0.05) between participant sexes (Cohen, 1988; Faul et al., 2007). In addition, we conservatively assumed the effect size of inversion as small-to-medium (Cohen’s *f* = 0.15) because of the mixed results of previous studies. A sample of 90 (45 for each group) is required to detect a significant difference (*α* = 0.05, 1 – *β* = 0.95, *r* = 0.7) between upright and inverted faces with a small-to-medium effect size, and a sample of 90 (45 for each group) is required to detect a significant interaction (*α* = 0.05, 1 – *β* = 0.95, *r* = 0.7) between the face orientation and sex with a small-to-medium effect size. Therefore, we decided to recruit a sample of more than 200 participants to confirm small-to-medium-sized effects between conditions and to overcome the potential loss of analyzable data, often caused by participants’ inappropriate responses or technical errors.

Two hundred and thirteen young adults participated. All had normal or corrected-to-normal vision, and none reported a history of neurological or developmental disorders. The study was conducted in accordance with the Declaration of Helsinki and was approved by the Waseda University Ethics Committee. Each participant gave written informed consent before the experiment. Twenty-three participants were excluded because their proportion of ‘direct’ responses on 0° trials was below 60%. This criterion also served as a basic post-hoc screen to ensure participants were engaged with the task and could perform it at a minimal level of proficiency. After these exclusions, data from 190 participants (self-reported 92 females, 98 males; mean age: 21.3 years; range: 18-29 years) remained for analysis.

### 2.2 Stimuli

Face images were obtained from three male and three female models, looking either directly at the camera (0°) or at 10°, 20°, or 30° to the left or right, with their faces square to the camera, and were photographed by the authors (Matsuyoshi et al., 2014). Images were grey-scaled and subtended approximately 12° × 17° of the visual angle. The mean luminance was normalized across images. A mask stimulus was generated by grid-scrambling the average face stimuli.

### 2.3 Procedure

Participants viewed the stimuli on a 21-inch CRT monitor at a distance of 57 cm with their heads resting on a chinrest and judged whether the eyes were looking directly at them (Matsuyoshi et al., 2014). All stimuli were presented against a black background. Each trial began with an upright or inverted face stimulus (20 ms) followed by a brief blank screen (20, 40, or 60 ms), a visual mask (100 ms), and finally, a fixation cross (until response) (Fig. 1). Face images were either direct gaze (0°) or left- or right-averted gaze (10°, 20°, or 30°). The vertical positions of the eyes were aligned between upright and inverted faces and located along the vertical centre of the display. We used a variable interstimulus interval (ISI) between the face and the mask to control task difficulty. Each left or right angle was presented 12 times, except for 0° stimuli, which were presented 24 times, for each of the three ISIs for both face orientations, resulting in 576 trials in total (6 gaze angles [left or right 10-30°] × 3 ISIs × 2 face orientations × 12 trials + 1 gaze angle [0°] × 3 ISIs × 2 face orientations × 24 trials). The two face orientations were presented separately in four alternating blocks of 144 trials each, with the order counterbalanced across participants (i.e., either upright-inverted-upright-inverted or inverted-upright-inverted-upright). Gaze directions and ISIs were randomised across trials. Participants performed the main experiment after 30 practice trials.

**Figure 1.**
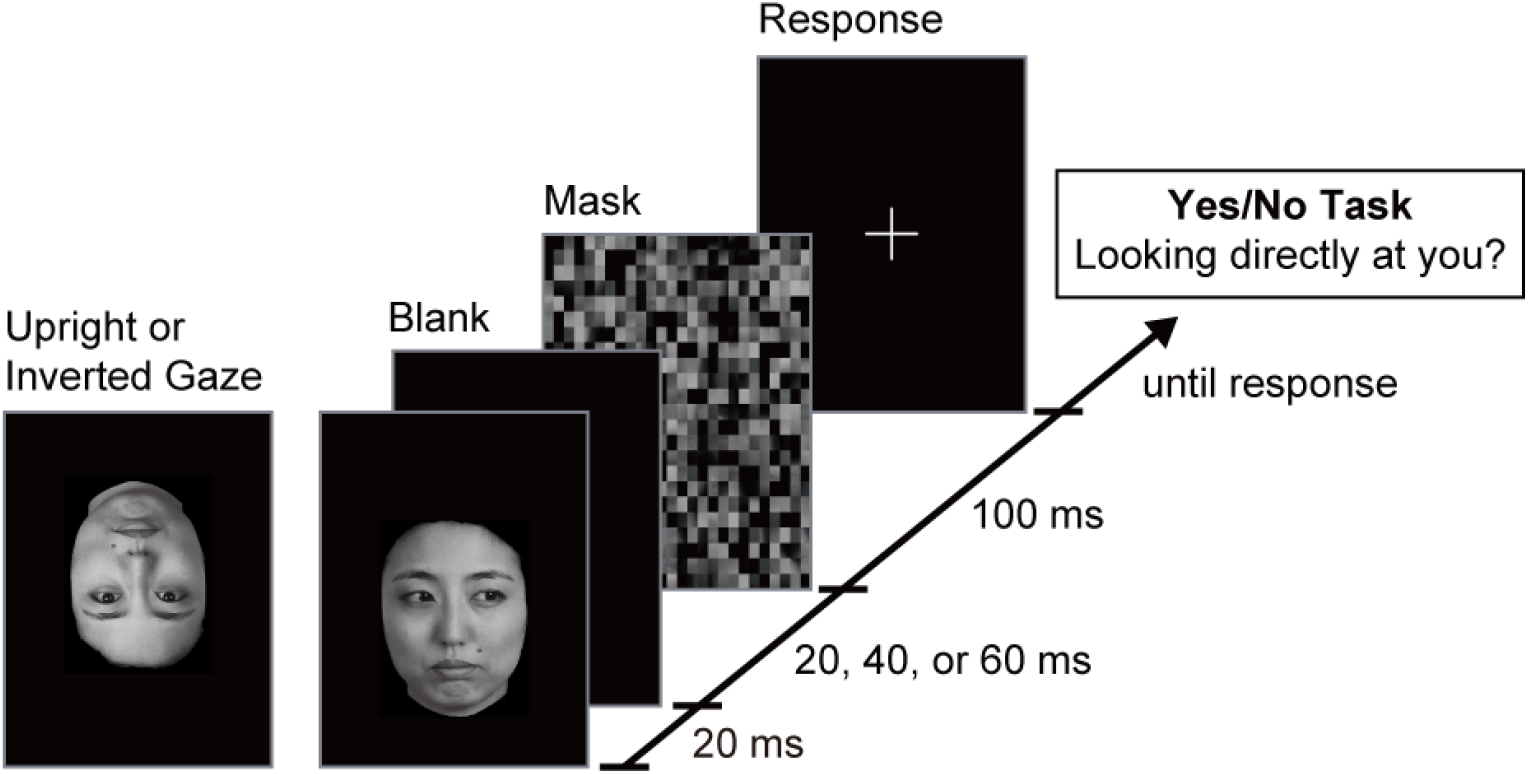
Experimental Paradigm. Schematic presentation of the experimental task. Upright and inverted faces were presented in different blocks (4 [2 upright + 2 inverted] blocks × 144 trials).

### 2.4 Analysis

Responses from left- and right-averted gaze stimuli were pooled for each absolute angle. In addition, responses from different ISIs were also pooled. The gaze threshold was defined as the angle at which a 50% response was obtained, as estimated by a local linear fit of a nonparametric psychometric function (Żychaluk & Foster, 2009) to each observer’s ‘direct’ responses as a function of gaze angle. This threshold represents the boundary of the ‘cone of direct gaze’ (i.e., the range of perceived direct gaze) (Gamer & Heiko, 2007; Jun et al., 2013), or the point of subjective equality for ‘direct’ vs. ‘averted’ judgments, rather than a discrimination threshold between two non-zero averted gaze angles as often used in typical psychophysical paradigms. A higher threshold indicated lower sensitivity to gaze direction. The significance level was set at α = 0.05.

## 3. Results

### 3.1 Psychometric functions

We performed a mixed ANOVA (with Greenhouse–Geisser correction applied where sphericity was violated) on the proportion of ‘direct’ responses, with angle (0°, 10°, 20°, 30°) and orientation (upright, inverted) as within-subject factors and participant sex (male, female) as a between-subject factor (Fig. 2a). The main effects of the angle (*F*_(1.99, 373.89)_ = 4796.912, *p* = 9.854 × 10^−270^, 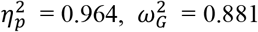, Cohen’s *f* = 2.737), face orientation (*F*_(1, 188)_ = 158.223, *p* = 1.007 × 10^−26^, 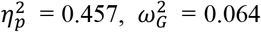, Cohen’s *f* = 0.091), and participant sex (*F*_(1, 188)_ = 9.758, *p* = 0.002, 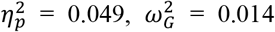, Cohen’s *f* = 0.044) were significant. This pattern indicates a decrease in ‘direct’ responses with increasing angle, higher ‘direct’ responses in the inverted compared to the upright face condition, and higher ‘direct’ responses in male compared to female participants, respectively.

**Figure 2.**
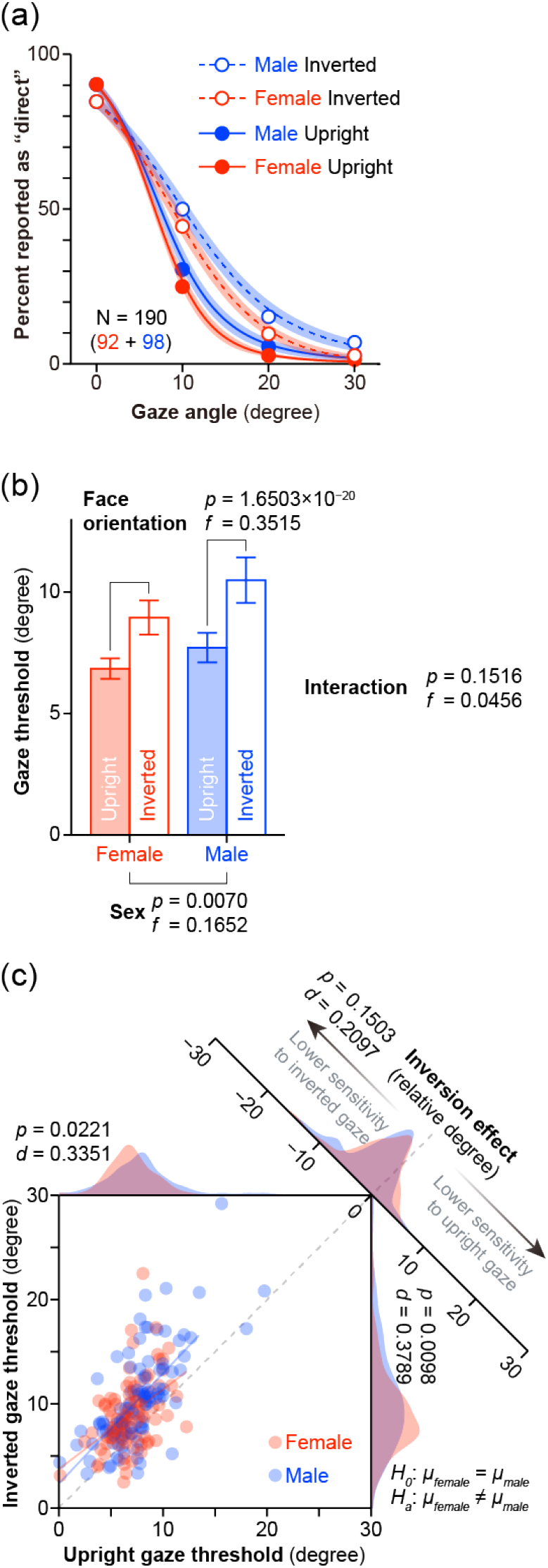
Experimental Results. (a) Gaze angle psychometric functions. Mean percentage of ‘direct’ responses (y-axis), plotted as a function of gaze angle (x-axis) × participant sex (female = red, male = blue) × face orientation (upright face = filled circle with a solid line, inverted face = open circle with dashed line). Shaded areas represent 95% confidence intervals of the mean. (b) Mean gaze threshold (angle of 50% reported as direct, which was estimated from the psychometric function) as a function of participant sex (female = red, male = blue) × face orientation (upright face = filled, inverted face = open). Error bars represent 95% confidence intervals of the mean. (c) Scatter plot and density curves of individual upright and inverted gaze thresholds (female = red, male = blue). The grey dashed line indicates identical gaze thresholds between upright and inverted faces (i.e., no gaze inversion effect).

We found significant interactions between the angle and orientation (*F*_(2.14, 402.85)_ = 276.829, *p* = 9.499 × 10^−80^, 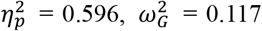, Cohen’s *f* = 0.127) and between the angle and participant sex (*F*_(1.99, 373.89)_ = 6.847, *p* = 0.001, 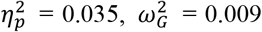, Cohen’s *f* = 0.035), while no significant interactions were found between the orientation and sex (*F*_(1, 188)_ = 1.622, *p* = 0.204, 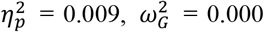, Cohen’s *f* = 0.009) and between the angle, orientation, and sex (*F*_(2.14, 402.85)_ = 0.962, *p* = 0.388, 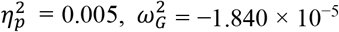, Cohen’s *f* = 0.007). Follow-up tests revealed higher ‘direct’ responses in the upright compared with the inverted face condition at an angle of 0° (*F*_(1, 188)_ = 68.290, *p* = 2.486 × 10^−14^, 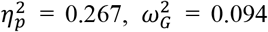, Cohen’s *f* = 0.325), and higher ‘direct’ responses in the inverted compared to the upright face condition at angles between 10 and 30° (*F*_(1, 188)_ values > 65.813, *p* values < 6.274 × 10^−14^, 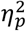 values > 0.259, 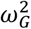 values > 0.084, Cohen’s *f* values > 0.298). In addition, male participants showed higher ‘direct’ responses at angles between 10 and 30° than female participants (*F*_(1, 188)_ values > 8.673, *p* values < 0.004, 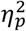 values > 0.044, 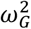 values > 0.031, Cohen’s *f* values > 0.165). By contrast, male and female participants showed similar ‘direct’ responses at an angle of 0° (*F*_(1, 188)_ = 0.215, *p* = 0.643, 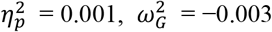, Cohen’s *f* = 0.027).

### 3.2 Gaze thresholds

A 2 (upright, inverted) × 2 (male, female) mixed ANOVA on the 50% gaze-perception threshold (Fig. 2b) showed a significant main effect of face orientation (*F*_(1, 188)_ = 109.666, *p* = 1.650 × 10^−20^, 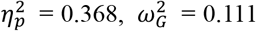, Cohen’s *f* = 0.352) with a larger threshold in the inverted compared to the upright face condition (medium-to-large effect size), and a significant main effect of sex (*F*_(1, 188)_ = 7.425, *p* = 0.007, 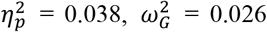, Cohen’s *f* = 0.165) with a larger threshold for male compared to female participants (small-to-medium effect size). However, there was no significant interaction between the face orientation and participant sex (*F*_(1, 188)_ = 2.073, *p* = 0.152, 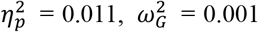, Cohen’s *f* = 0.046), showing that both sexes showed comparable inversion effects (females, *t*_(*91)*_ = 6.639, *p* = 2.244 × 10^−9^, Cohen’s *d* = 0.692 [95% CI: 0.394, 0.989]; males, *t*_(*97)*_ = 8.174, *p* = 1.142 × 10^−12^, Cohen’s *d* = 0.826 [95% CI: 0.533, 1.117]; females − males, *t*_(*187*.*75)*_ = 1.444, *p* = 0.150, Cohen’s *d* = 0.210 [95% CI: −0.076, 0.495]), whereas female participants had a higher sensitivity than male participants for discriminating gaze direction irrespective of face orientation (upright, *t*_(*170*.*18)*_ = 2.309, *p* = 0.022, Cohen’s *d* = 0.335 [95% CI: 0.048, 0.621]; inverted, *t*_(*177*.*05)*_ = 2.6104, *p* = 0.010, Cohen’s *d* = 0.379 [95% CI: 0.091, 0.666]).

There was a significant correlation between the upright and inverted gaze thresholds (Fig. 2c; overall, *r* = 0.631, *p* = 1.556 × 10^−22^; males, *r* = 0.693, *p* = 2.531 × 10^−15^; females, *r* = 0.461, *p* = 3.844 × 10^−6^), indicating that there are partly shared mechanisms in upright and inverted gaze processing. The correlation was significantly greater in male than in female participants (*z* = 2.415, *p* = 0.015). Note, however, that the difference was not robust to the exclusion of two outliers (one male and one female participant with the largest inverted gaze thresholds), after which it was no longer significant (*z* = 1.883, *p* = 0.060).

## 4. Discussion

Using a well-powered cohort with an a priori determined sample size, the present study showed that inverting the face reliably impaired sensitivity to small deviations from direct gaze. At the same time, we observed higher overall gaze sensitivity in females than in males, while the magnitude of the inversion cost was closely comparable across sexes. We recognise that gaze perception involves distinctive signalling properties, such as the high contrast between the iris and sclera, which may set it apart from other facial information (Kano, 2023; Kobayashi & Kohshima, 1997). It would therefore be inappropriate to assume that the present results generalise automatically to all facial features or that they fully resolve the broader debate between qualitative and quantitative views of inversion effects. Rather, they provide a useful boundary condition for these theories. Specifically, they are difficult to reconcile with a strict qualitative view that treats feature-level processing as largely preserved under inversion. The fact that inversion produces a clear decrement in performance for a task that can in principle be supported by relatively local eye information suggests that this assumption does not hold for at least one highly salient facial cue (eye gaze). Our data do not establish that all feature processing is governed by purely quantitative differences, but they do show that the qualitative prediction of preserved featural processing cannot be universal.

### 4.1 Qualitative and quantitative views of the inversion effects

Several studies have suggested that inversion preserves featural processing while severely impairing holistic or configural processing, such as the distances between features (Freire et al., 2000; Le Grand et al., 2001; Rossion, 2008; Tanaka & Sengco, 1997; Young et al., 1987). On this qualitative view, inversion is assumed to prompt a shift toward feature-based encoding, so that performance on tasks driven by individual features should be relatively similar for upright and inverted faces (Farah et al., 1995). In particular, the eyes have sometimes been regarded as the least susceptible feature to inversion. For example, Xu and Tanaka (2013) reported that inversion did not affect the perception of changes in the eye region, whereas it degraded sensitivity to changes in the mouth region (N = 22, 40% female). Although their finding appears inconsistent with ours, their task presented faces for 500 ms, which is substantially longer than the 20-ms exposure used here. Such long encoding durations may have attenuated or masked any disadvantage for inverted relative to upright faces, and behavioural performance in their study was near ceiling (about 90%), leaving limited room for inversion costs to emerge.

By contrast, quantitative views posit that similar mechanisms are engaged for upright and inverted faces but operate less efficiently when faces are inverted (Gold et al., 2012; Jiang et al., 2006; Sekuler et al., 2004). Jenkins and Langton (2003) reported that inversion impaired sensitivity to gaze direction, although their study used a between-participants design, a small sample size (N = 6 per group), and an unknown sex ratio. Furthermore, Rakover and Teucher (1997) (N = 16, 73% female) and Leder et al. (2001) (N = 20, 90% female) likewise found it more difficult to process individual features when they were inverted. However, the limited and often unbalanced samples in these studies make it difficult to draw firm conclusions about the size and robustness of the effect. Our findings help clarify the extent to which inversion affects gaze perception in a large, sex-balanced sample, using tightly controlled psychophysical methods.

Although prior evidence hinted that inversion can impair gaze direction sensitivity (Jenkins & Langton, 2003), the strength and generality of this effect had not been firmly established across paradigms and larger cohorts, leading to ambiguity in the literature. The present study was designed to address this gap by quantifying the gaze inversion cost with greater precision in a well-powered cohort with a balanced sex ratio.

It is also important to recognise that the questions of ‘qualitative versus quantitative’ and ‘configural versus featural’ are conceptually distinct, even though they are intertwined in empirical work (Gold et al., 2012; Maurer et al., 2002; Sekuler et al., 2004). The qualitative view typically links upright faces with configural processing and inverted faces with featural processing. Quantitative accounts, by contrast, do not assume such an orientation-dependent processing style; instead, they propose that inversion generally reduces processing efficiency. Our design targeted a key implication of a *strict* qualitative account: if featural processing is preserved or even enhanced for inverted faces, then performance on a task based largely on information from the eye region should show little or no impairment under inversion. In contrast to this prediction, we observed a clear impairment in gaze discrimination when faces were inverted. This suggests that, at least for gaze judgements of the sort studied here, inversion costs are not confined to clearly global, configural aspects of faces but extend to tasks that can be supported largely by eye-region cues.

### 4.2 The effects of task settings on inversion effects

Although we demonstrated an inversion-induced impairment in gaze sensitivity, many other studies (Freire et al., 2000; Hole, 1994; Le Grand et al., 2001; Young et al., 1987), including large-sample investigations (N = 242–282) (Rezlescu et al., 2017; Susilo et al., 2013), have reported preserved or even improved performance when comparing individual features in inverted faces. These apparent discrepancies are unlikely to result from low statistical power or an imbalanced sex ratio, since both these large-sample studies and the present one used sizeable cohorts, and we (48% female) and Susilo et al. (2013) (52% female) achieved approximately balanced sex ratios. We suggest instead that task settings may be a major source of divergence. Featural processing is sometimes preserved (or enhanced) despite the diminished sensitivity, perhaps not because parts-based processing is dominant for inverted faces (Farah et al., 1995; Tanaka & Farah, 1993), but because the tasks require discrimination between exemplars from the same facial parts (e.g., comparing the eyes of different people). Thus, even though perceptual sensitivity per se decreases, the ability to compare two signals may be relatively unlikely to deteriorate because sensitivity to both signals, but not to either, decreases. Moreover, the apparent dominance of feature-based processing for inverted faces may result from greater difficulty decomposing facial parts in upright faces, rather than from enhanced featural processing for inverted faces. Processing an upright face likely involves tight integration of internal features, making it difficult to parse the face into isolated features (Maurer et al., 2002); however, an inverted face is less likely to undergo this integration. Under such circumstances, explicit behavioural performance may depend less on whether inversion alters the underlying processing style and more on how the specific task elicits comparisons between stimuli, the distribution of attention across features, and the overall processing load. In this context, our finding that inversion impairs gaze sensitivity in a task that emphasises brief, single-stimulus judgements complements prior work using comparison tasks and suggests that different paradigms may reveal different facets of how inversion affects feature-level information.

Note that the effect size of the GIE was smaller than that of the typical FIE. Despite a huge effect of the FIE in a face recognition task (*d* = 1.298) (Matsuyoshi et al., 2015), we found a medium-to-large effect of the GIE in the present study (*d* = 0.761, across all participants). One reason for the tremendous difference in effect size might be that a whole face accumulates the impairments in the processing of multiple facial features (Gold et al., 2012). Although susceptibility to inversion may differ across parts (Rakover & Teucher, 1997; Xu & Tanaka, 2013), these differences may aggregate to produce a large effect size for the FIE. Alternatively, the large effect size difference may simply reflect differences in processing load between the tasks. The present task encouraged observers to focus on the eyes. By contrast, typical FIE tasks do not explicitly encourage observers to focus on a facial feature but rather on a whole face. Encoding a whole face may be more challenging than encoding a single facial feature.

### 4.3 Sex differences in face and gaze processing

Consistent with numerous studies reporting female advantages in face processing (Bayliss et al., 2005; Goodman et al., 2012; Heisz et al., 2013; Matsuyoshi et al., 2014; Matsuyoshi & Watanabe, 2021; McClure, 2000), we found that females outperformed males in gaze direction sensitivity. The effect size for this female advantage in upright gaze sensitivity (Cohen’s *d* = 0.3351) was notably smaller than that reported for upright face recognition in a previous study from our group (Cohen’s *d* = 0.643, N = 180, 45% female) (Matsuyoshi & Watanabe, 2021). This may indicate that tasks that rely more on relatively local feature information, such as brief gaze judgements, are likely to manifest weaker effects than whole-face processing, perhaps because this task draws primarily on a single feature, so inversion-related costs may not compound across multiple features.

The observed female superiority in gaze sensitivity may reflect that females use more efficient scanning strategies or have a lower threshold for detecting subtle social signals. While the exact origin of this difference remains debated, prior work suggests that both biological factors (Bölte et al., 2023; Kiesow et al., 2020; Kundakovic & Tickerhoof, 2024) and psychological factors, such as internalized social norms or stereotypes (Briton & Hall, 1995; Crandall et al., 2002; Gavrilets & Richerson, 2017; Rutland et al., 2005), may contribute to sex and gender differences in social cognition. In the present study, females showed higher baseline sensitivity, perhaps reflecting some combination of these factors, but the relative cost of inversion was similar for females and males. This pattern suggests that the computations supporting gaze discrimination are similarly affected by inversion across sexes, even though baseline performance levels differ.

A notable limitation of our study is that we treated sex/gender in a binary fashion, and findings solely depended on self-reported sex/gender. Outcomes can vary across individuals with different birth-assigned sexes and gender identities (Bölte et al., 2023; Joel, 2021). Definitions of sex/gender are continually evolving, and sex/gender is now understood as a spectrum rather than as simple binary categories (Ainsworth, 2015; Kundakovic & Tickerhoof, 2024; Rosenthal, 2021). Future research that more fully incorporates variation across the sex/gender spectrum will be necessary for understanding how biological and sociocultural factors jointly shape face and gaze processing.

### 4.4 Conclusion

In conclusion, our study provides robust evidence for an inversion-induced impairment in gaze direction sensitivity (GIE). While our findings do not resolve the broader debate over the nature of the FIE, they directly challenge a strict interpretation of the qualitative view, which posits that the processing of individual facial features is preserved under inversion. Thus, they nonetheless provide a critical boundary condition for theories assuming preserved parts-based processing. Our results demonstrate that, at least for the crucial feature of eye gaze, sensitivity is significantly diminished when the face is inverted, even in a task that can in principle be supported by information within the eye region. Since we did not incorporate tasks other than discriminating gaze direction, further research is required to clarify the conditions under which processing of facial feature information is preserved or impaired by inversion. Nevertheless, our findings establish the GIE as a reliable phenomenon for brief gaze judgements and highlight the value of using well-powered, sex-balanced samples when investigating how inversion shapes both face perception and individual differences in gaze processing.

## Acknowledgements

This study was supported by grants from the Japan Society for the Promotion of Science (JP19H04433 to DM; JP22H00090 and JP25H01234 to KW). The photographed model acknowledges the use of the face images in Figure 1 with written informed consent. The dataset analyzed during the current study is available at GitHub:https://github.com/dicemt/matsuyoshi_gaze_inversion_effect

## Abbreviation

FIE: face inversion effect
GIE: gaze inversion effect

## Authors’ contributions

All authors designed the study and wrote the manuscript. DM and KK collected stimulus materials. DM collected and analyzed the data.

